# Low amplitude burst detection of catecholamines

**DOI:** 10.1101/2021.08.02.454747

**Authors:** Amnah Eltahir, Jason White, Terry Lohrenz, P. Read Montague

**Affiliations:** Fralin Biomedical Research Institute at VTC, Virginia Tech, Roanoke, VA 24016, USA; Department of Physics, Virginia Tech, Blacksburg, VA 24061, USA

## Abstract

Machine learning advances in electrochemical detection have recently produced sub-second and concurrent detection of dopamine and serotonin during perception and action tasks in conscious humans. Here, we present a new machine learning approach to sub-second, concurrent separation of dopamine, norepinephrine, and serotonin. The method exploits a low amplitude burst protocol for the controlled voltage waveform and we demonstrate its efficacy by showing how it separates dopamine-induced signals from norepinephrine induced signals. Previous efforts to deploy electrochemical detection of dopamine *in vivo* have not separated the dopamine-dependent signal from a norepinephrine-dependent signal. Consequently, this new method can provide new insights into concurrent signaling by these two important neuromodulators.

## Introduction

There exist many techniques to probe physiological changes in the brain in response to cognitive demands, including functional magnetic resonance imaging (Ogawa & Lee, 1990), electroencephalography (Behrens et al., 1994) and magnetoencephalography (Hämäläinen, Hari, Ilmoniemi, Knuutila, & Lounasmaa, 1993). Nevertheless, there persists a gap in the ability to track sub-second chemical changes in human brain related to synaptic transmission (Kandel & Siegelbaum, 1995). Techniques such as positron emission tomography (Wrenn, Good, & Handler, 1951) and single emission computed tomography (Takeshita et al., 1992) map the activity of neurotransmitter receptors; however, their limited temporal resolution makes them unsuitable for the study of sub-second neural events (Dayan, 2012). Recent work adapting fast scan cyclic voltammetry (FSCV), (Kissinger, Hart, & Adams, 1973; Kuhr & Wightman, 1986; Stamford, Kruk, & Millar, 1986) from animal experiments to clinical settings in humans has opened a new field of invasive real-time neurochemistry (Bang et al., 2020; Kishida et al., 2016; Montague & Kishida, 2018; Moran et al., 2018).

In FSCV studies, experimenters implant an electrode into the cell cultures of brain tissue (Adams, Puchades, & Ewing, 2008), as well as brains of a living animal, including rodents (Rebec, 1998), primates (Schluter, Mitz, Cheer, & Averbeck, 2014), and zebrafish (Jones, McCutcheon, Young, & Norton, 2015). A voltage of a specified waveform is applied to the electrode, producing electrical current responses on the surface of the electrode. The current traces can then be used to make inferences about the identity and relative concentration of oxidizable neurotransmitters like dopamine, serotonin, and so on (Kissinger & Heineman, 1983). In the case of dopamine, a voltage sweep produces oxidation and reduction peaks at characteristic voltage potentials which can be calibrated ex vivo to known changes in concentration (Heien, Johnson, & Wightman, 2004). Such an invasive method does not translate readily into humans because the opportunity to pre-calibrate the electrode is not available. However, surgeries which involve electrode implantation, such as deep brain stimulation (DBS) implantation surgery, provide an exciting opportunity to obtain electrochemical recordings from conscious patients performing a task (Bang et al., 2020; Kishida et al., 2016; Moran et al., 2018).

The process of adapting FSCV methods for human use has revealed that catecholamines like dopamine show small but significant oxidation through a wide range of applied voltages (Kishida et al., 2011; 2016; Moran et al., 2018; Bang et al., 2020 shows datasets used for training models). One change in approach has been to use all the current time-series data throughout the duty cycle of an applied voltage waveform, and then deploy various machine learning methods with proper cross-validation to produce excellent concentration prediction models for dopamine and serotonin (Moran et al., 2018; Bang et al., 2020). To date, these models have employed elastic net (EN)-penalized regression, a common machine learning algorithm (Friedman, Hastie, & Tibshirani, 2010; Zou & Hastie, 2005).

Here, we develop and validate a voltage waveform at a significantly reduced voltage range relative to standard FSCV protocols. The voltage range used by older methods to measure dopamine spans -0.6 V to 1.4 V, in order to capture oxidation and reduction peaks. Devising a low amplitude sensing method that remains sensitive to changes in dopamine concentration would remove the restriction on potential range – a significant advance when considering its use in human subjects. The low amplitude sensing method must also identify multiple (possibly confounding) neurotransmitters, as would be the case in-vivo. This includes detecting neuromodulators with similar electrochemical responses, such as the catecholamines dopamine and norepinephrine. Lastly, the low amplitude method must work on data gathered on probes not used during the model generating process, as would be the case for probes used in subjects. That is, we seek models that generalize both out-of-concentration and out-of-probe.

The first experiment explored two different voltage waveforms at one quarter of the range used in fast scan cyclic voltammetry (FSCV) protocols (Rodeberg, Sandberg, Johnson, Phillips, & Wightman, 2017). Traditional FSCV makes use of characteristic oxidation peaks to calibrate readings and estimate changes in neurotransmitter concentration (Baur, Kristensen, May, Wiedemann, & Wightman, 1988). The first voltage waveform used a triangular pulse, with a shape similar to FSCV (Howell, Kuhr, Ensman, & Mark Wightman, 1986), sweeps but limited to -0.15 to 0.35 volts. The second used a randomly ordered voltage waveform also limited to -0.15 to 0.35 volts (this is analogous to waveform discussed in Montague et al., 2019). The second experiment demonstrates the capacity of this low amplitude technique to separate signals due to dopamine from those due to norepinephrine.

## Results

### Comparing low amplitude forcing functions

Recordings were obtained on dopamine solutions ranging from 0-2.7 µM in steps of 50 nM with added Gaussian jitter, and within-probe models outlined in the methods section. The voltage waveforms used were the modified low amplitude FSCV in Fig. 1A and low amplitude random burst in Fig. 1B, with an average action potential plotted in red for scale. These voltage waveforms were delivered in voltage clamp mode and the current time series were recorded. We used the time derivative of the current time series to predict the known dopamine concentration – this is a standard supervised learning problem. The ‘reported oxidation potential for dopamine’ is around +0.65 volts so this triangular waveform peak is well shy of that reported potential. Nevertheless, there is dopamine oxidation and reduction throughout the waveform, and we pursued the hypothesis that many machine learning methods could pick out predictive information We used the elastic net algorithm (Zhou and Hastie, 2004) to build a cross-validated concentration prediction model (Kishida et al., 2016; Moran et al., 2018; Bang et al., 2020). Within-probe models generated predictions which are plotted in Fig. 1C for the low amplitude FSCV and Fig. 1D for the low amplitude random burst waveforms. The dopamine predictions in blue are plotted over the true values in black found in the top panel. The dopamine predictions show a closer fit and reduced noise in prediction for the low amplitude random burst. These findings are quantified in the lower left panel for RMSE and lower right panel showing SNR. The values are plotted for each concentration, along with the average as a dotted line. Overall, the low amplitude random burst sensing demonstrates lower RMSE and higher SNR than the low amplitude sweep. One known contributing adulterant to a good model in this context is autocorrelation, which is largely removed by the randomized sequence of voltages (Montague et al., 2019).

**Figure 1.**
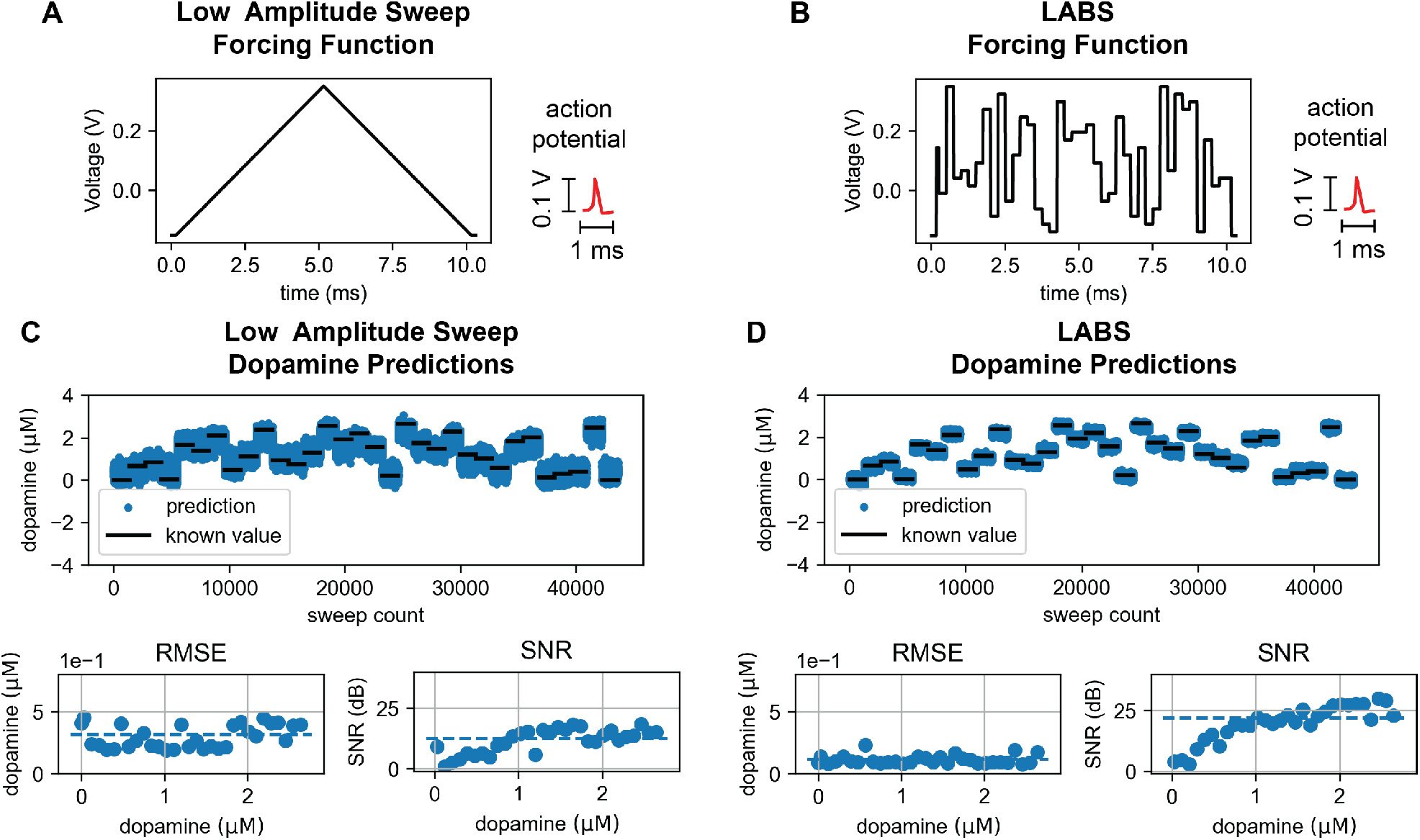
Low amplitude linear sweep versus random burst. (A) A low amplitude version of the standard sweep protocol, with action potential plotted in red for scale. (B) The LABS forcing function, with action potential plotted in red for scale. (C) Dopamine predictions using low amplitude sweep plotted over known values, with the RMSE in the lower left panel and the SNR in the lower right panel. (D) Dopamine predictions using low amplitude random burst sensing plotted over known values, with the RMSE in the lower left panel and the SNR in the lower right panel.

### Distinguishing dopamine and norepinephrine

Using the random burst voltage protocol, we sought to separate dopamine and norepinephrine predictions. In all previous work estimating dopamine using FSCV, it has not been possible to accomplish such a separation because of their extremely similar current profile near the reported oxidation points for both. EN-penalized linear regression models were generated using the within-probe modeling procedure outlined in to calibrate both dopamine and norepinephrine concentrations. The voltage waveform for FSCV plotted in Fig. 2A shows how the function LABS waveform spans a voltage potential range well below the oxidation peak for dopamine. In Figs. 2B and 2C, the results of EN-penalized regression models are plotted against the known values for both methods, showing a strong direct relationship with a slope greater than 0.99. The correlation between the known and predicted concentrations, indicated by correlation coefficients R^2^ greater than 0.99 for both neurotransmitters.

**Figure 2.**
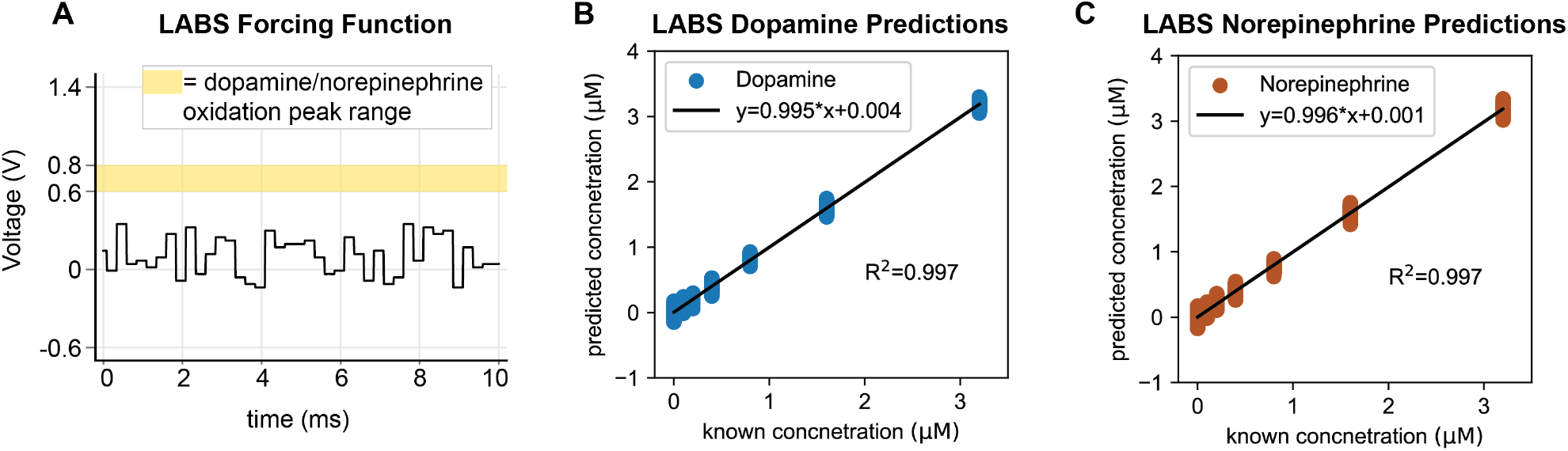
Distinguishing dopamine and norepinephrine. (A) LABS forcing function relative to oxidation peak region of dopamine and norepinephrine. (B) Within-probe EN-penalized model predictions for dopamine versus known concentrations. The line of best fit is reported in the legend, and the accuracy is reported in terms of R^2^. (C) Within-probe EN-penalized model predictions for norepinephrine versus known concentrations. The line of best fit is reported in the legend, and the accuracy is reported in terms of R^2^.

### Out-of-probe predictions

Low amplitude random burst sensing data were acquired on ten probes using randomized mixtures of dopamine and serotonin from 0-2.7 µM in steps of 50 nM with randomized pH, ranging from 6.8 to 7.8. An ensemble model approach, outlined in the methods and Fig. 4C, was built to predict each probe, in which data from the probe to be tested was excluded from model generation. The training data from the remaining probes was used to build an ensemble model. The testing data were similarly partitioned, and the EN-model from within the ensemble closest to the average predictions was selected. An example for results for a single probe in Fig. 3A shows a direct relationship between the known and true values of neurotransmitters, with R^2^ close to 0.9 for dopamine and over 0.94 for serotonin, indicating the ensemble model captured the variance in neurotransmitter changes across the different solutions collected on this probe. The model also captured changes in pH, with an R^2^above 0.79, indicating a strong effect size.

**Figure 3.**
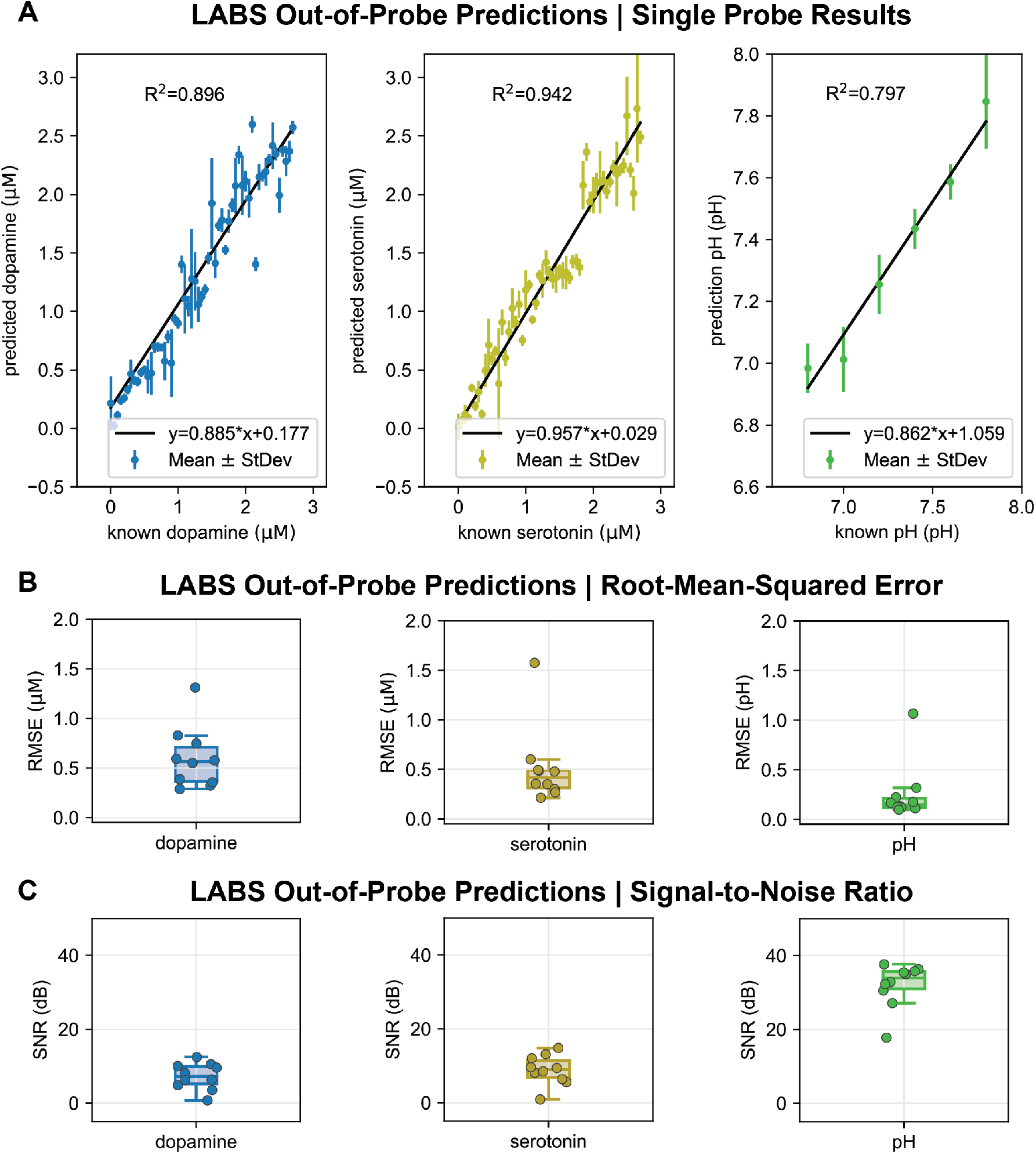
LABS out-of-probe predictions of analytes. (A) An example of predictions for dopamine (green) serotonin (yellow), and pH (green) plotted against known values, along with the line of best fit. The predictions are plotted as mean and standard deviation at each concentration and pH. Each legend contains the equation for line of best fit between known and predicted values. The R^2^ for dopamine is close to 0.9, greater than 0.9 for serotonin, and greater than (B) Box plots of average RMSE for dopamine, serotonin and pH for all ten probes. (C) Box plot of the average SNR for dopamine, serotonin and pH for all ten probes.

**Figure 4.**
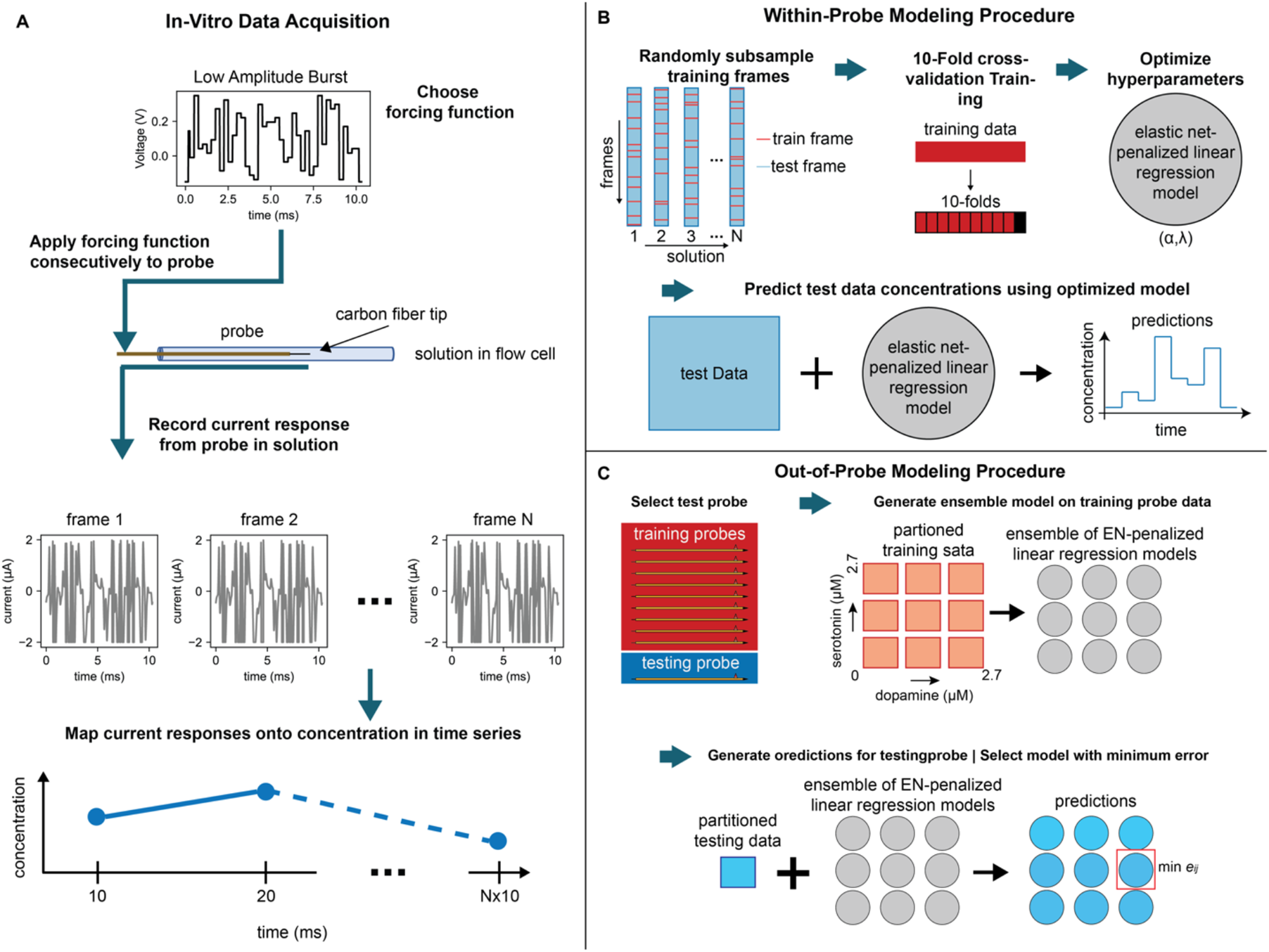
Graphical Depiction of data collection and modeling procedures. (A) In-vitro data collection starts by applying a specified forcing function through a probe submerged in solution. The current “frames” produce neurotransmitter concentration estimates, which arranged in sequential order to provide a time series of concentrations. (B) Within-probe modeling procedure starts by constructing a training set from randomly subsampled sweeps. Ten-fold cross-validation training using EN-penalized regression produces a linear model, parameterized over *α* and *λ*. The model then predicts concentrations from independent test data. (C) Out-of-probe modeling procedure leaves out one probe to test and trains a model on the remaining ten probes. The data are subdivided into bins based on concentration of dopamine and serotonin, for which each will generate an EN-penalized linear regression model. The testing data were similarly partitioned and each of the models in the ensemble produce a set of predictions. The distance e_ij_, defined in Eq. 6, is used to select the model which lies closest to the average of the predictions.

Statistical summaries of all ten probes show an average RMSE below one µM in both dopamine and norepinephrine, for the exception of one outlier. While not directly comparable, pH has an average RMSE below 0.5, except for one outlier. The outlier may indicate data quality issues for a single probe within the collection of probes. The average SNR for all ten probes were near 10 dB for dopamine and norepinephrine. Dopamine and serotonin have similar distributions SNR values, while the distribution SNR of pH lies much higher.

## Discussion

The capabilities of electrochemical sensing at low voltage amplitudes were explored using two waveforms. The results of comparing dopamine predictions at the same low amplitude using both a linear sweep and burst forcing function indicates the random burst provides more accurate results at reduced amplitudes. A potential reason for the difference in model performance could be the relatively diminished autocorrelation of a random burst function pairs well with an algorithm like EN-penalized linear regression, a method that best suited for sparse signals (Candes & Wakin, 2008). There remain numerous untested possibilities for optimizing forcing function design.

The ability to distinguish dopamine and norepinephrine using LABS demonstrates an important step forward in electrochemical sensing. The high prediction accuracy of the models demonstrates that information content needed to track neurotransmitters exists outside the peak oxidation potential of the target neurotransmitters, validating the assertion that electrochemical protocols need not span the specified voltage range of FSCV. Differentiating neurotransmitters with similar electrochemical profiles without additional pharmacological aid or neural stimulation allows invasive neurochemical recordings to produce chemical specificity.

Proving the ability of LABS to make predictions out-of-probe is crucial to demonstrating feasibility of this method in-vivo. The results of the out of probe models to predict dopamine, serotonin and pH in mixture for a collection of ten probes demonstrate the ability to identify changes in multiple analytes changing simultaneously. For the exception of one outlier, all out-of-probe model predictions had a RMSE for dopamine and serotonin in mixture below one µM, which has yet to be demonstrated with any other modality at this temporal resolution. The outlined ensemble method demonstrates just one of many possibilities for building robust, chemically specific out-of-probe models to apply to in-vivo data.

Low amplitude burst sensing arose from the need to adapt a decades old voltammetry methodology for human populations. The reduction in applied voltages, temporal resolution and chemical specificity indicate there remain unexplored possibilities in electrochemistry to optimize protocols for biological applications. With the feasibility of LABS as sensing method for tracking multiple neurotransmitters has established, further testing can proceed to employ this method in-vivo, as well as further in-vitro testing to determine optimized sensing protocols.

## Methods

### In-vitro preparation

Voltammetry recordings were collected on in-vitro solutions of neurotransmitters, including dopamine, norepinephrine and serotonin, diluted to predetermined concentrations in phosphate buffered saline (PBS) solution. The neurotransmitters in powdered form were initially diluted in 0.1 N HCl to 100 mM stock solutions. The neurotransmitters were further diluted in aliquots containing 6.8 pH PBS before being prepared according to a predetermined solution schedule. Solutions were loaded into syringes and pushed into cylindrical glass flow cells that held the working end of the electrode. To limit time related signal drift, solution schedules were randomized. The solutions in the first experimental results section comparing low amplitude waveforms consisted of 7.8 pH PBS solutions with dopamine concentrations ranging for 0-2.7 µM in steps of 50 nM, with variation in steps added as gaussian noise. The solutions in the second experimental results section consisted of dopamine and norepinephrine at concentrations 0, 0.2, 0.4, 0.8 1.6 and 3.2 µM in solutions of 7.4 pH PBS. Next, results in Solutions in the final experimental results section consisted of data collected on ten probes used to create out-of-sample models were acquired on PBS solutions 0-2.7 µM concentrations in steps of 50 nM of dopamine and norepinephrine, both separately and in mixture, at PBS ranging in pH from 6.8 to 7.8. Data sets collected on different days had different randomizations of solutions and selections of mixtures.

### Probe construction

The probes used to collect voltammetry data were constructed in the laboratory based on FSCV carbon-fiber electrodes in the literature (Kishida et al., 2016). The probe manufacturing process began by threading a 1.2 cm length, 7 um diameter, carbon fiber into a (reference no. LS330423: Goodfellow) into a one centimeter fused one-centimeter-long silica capillary coated with biocompatible polyimide coating (Polymicro Technologies). The carbon fiber was affixed to the assembly by pulling it through a droplet of epoxy placed at the end of the silica tubing. After drying overnight, the tip was trimmed to one millimeter beyond the edge of the assembly to form the working tip of the electrode. Another assembly was constructed by threading 29 cm long platinum iridium wire through a 28 cm long polyimide-coated capillary (Polymicro Technologies). The two assemblies were combined using silver paint and dried over 24 hours. Gold pins connectors used to hook up to the headstage were soldered onto the non-working probe end. The entire fabrication was then placed inside of a stainless-steel guide tube of similar construction to those used in DBS-electrode implantation surgery and secured at the microsensor assembly using two-part epoxy such that the working tip protruded one centimeter from the end of the guide tube. For more detailed explanations of methods probe construction procedures, see previous works (Kishida et al., 2016, 2011).

### Data acquisition and preprocessing

Voltammetry data were acquired using an electrophysiology system consisting of a head stage (CV-7B/EC; Axon Instruments), amplifier (700B; Axon Instruments; Multiclamp), analog-to-digital converter (Digidata; Axon Instruments) and laptop (MacBookPro; Apple). Using pClamp controller software, the sampling frequency was set to 100 kHz. The upper limit of 10,000 frames per file were collected per solution when using 97 Hz low amplitude protocols and 1000 frames for the 10 Hz FSCV protocol. Prior to each round of data collection, 30 seconds of pre-cycling using a truncated 10 millisecond FSCV forcing function were used to equilibrate the signal. The workflow in Fig. 4A shows the how applied forcing functions produce current “frames” which correspond to known concentrations of neurotransmitters. Recorded frames were saved as axon binary files (abf) for each solution.

To ensure quality and reproducibility of machine learning models, data were preprocessed using standard Python methods and libraries. Recorded frames of data stored in the raw abf files were truncated by removing the first and last 16 data points, leaving one thousand frame points per frame. In lieu of background subtraction, the frames were differentiated along the index, yielding data with 999 features per frame used as inputs in machine learning models. For each set of recordings, a stable window of frames – defined by the median current frame value – was identified and extracted. For each set of experiments, a stable 1500 frame window was selected based. Outliers, defined by median absolute deviations, were removed from each frame window. Within each window, 125 frames were randomly subsampled to train EN-penalized linear regression models. The remaining frames were used as testing data to evaluate the model accuracy.

### EN-Penalized Linear Regression Calibration

This study followed recent trends in human voltammetry by using EN-penalized linear regression to calibrate voltammetry (Bang et al., 2020; Kishida et al., 2016; Moran et al., 2018). The preprocessed in-vitro training data act as the input, while the concentrations and pH act as training in the regression analysis performed by the Python package GLMnet (Balakumar, 2016; Zou & Hastie, 2005). The regression coefficients in β correspond to each point along the differentiated frame. EN-penalized regression applies a penalty term defined by two hyperparameters, α and λ, which constrain the sum of squares residual to be minimized, as described in Eq. 1.

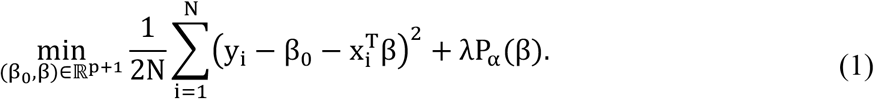

The penalty term P_*α*_(β), described in Eq. 2, is defined by its size, λ, while the ratio between *ℓ*_1_ and *ℓ*_2;_ norms, α, which ranges from zero to one.

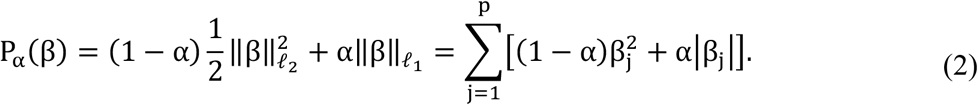

### Within-probe models

Preprocessed training data were used as inputs in EN-penalized regression training, as outlined in the workflow in Fig. 4B. Randomized ten-fold cross validation was carried out to optimize λ for α from zero to one in steps of 0.1. The combination of hyperparameters with minimum cross-validation error were used to optimize the linear regression model. Within-probe models were used to calibrate training data. Predictions generated on test data were used to evaluate model accuracy in terms of root-mean-squared error (RMSE) and signal-to-noise ratio (SNR).

### Out-of-probe models

Out-of-probe models, or models generated from probes independent of the one being tested, were created using an ensemble method outlined in Fig. 4C. Ensemble models were generated based on concentration bin, with each probe in the collection of probes left out for testing, w The data were partitioned into bins defined by concentration ranges, with dimensions of 900 nM dopamine by 900 nM serotonin, as shown in Table 1. Model training for each bin followed the same ten-fold cross-validation EN-penalized regression like the one described in the “Within-probe models” section. The testing data were similarly partitioned according to the same concentration bins. The reasoning behind restricting the concentration range of data lies in the assumption that concentrations of neurotransmitters would not fluctuate dramatically in-vivo. For each partitioned test data set, all ensemble model regressions were used to predict dopamine, serotonin and pH. The optimal model for each set of test data according to the distances e_i,j,DA_ and e_i,j,5HT_ between dopamine and serotonin predictions and the middle of the bin, defined in Eqns. 4 and 5, and choosing the bin with minimum combined distance e_i,j_, defined in Eq. 6.

**Table 1.** Ensemble model data configuration: Data bins defined by concentrations of dopamine and serotonin used to generate ensemble model for out-of-probe predictions.

|  |  |  |  |
| --- | --- | --- | --- |
| $r \quad j \rightarrow$ | | | |
| $i \quad \downarrow$ | DA: [0, 900] nM<br>5HT: [0, 900] nM<br>pH: [6.8, 7.8] | DA: [0, 900] nM<br>5HT: (900, 1800] nM<br>pH: [6.8, 7.8] | DA: [0, 900] nM<br>5HT: (1800, 2700] nM<br>pH: [6.8, 7.8] |
|  | DA: (900, 1800] nM<br>5HT: [0, 900] nM<br>pH: [6.8, 7.8] | DA: (900, 1800] nM<br>5HT: (900, 1800] nM<br>pH: [6.8, 7.8] | DA: (900, 1800] nM<br>5HT: (1800, 2700] nM<br>pH: [6.8, 7.8] |
|  | DA: (1800, 2700] nM<br>5HT: [0, 900] nM<br>pH: [6.8, 7.8] | DA: (1800, 2700] nM<br>5HT: (900, 1800] nM<br>pH: [6.8, 7.8] | DA: (1800, 2700] nM<br>5HT: (1800, 2700] nM<br>pH: [6.8, 7.8] |

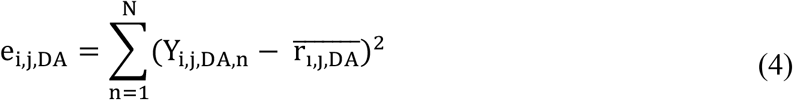

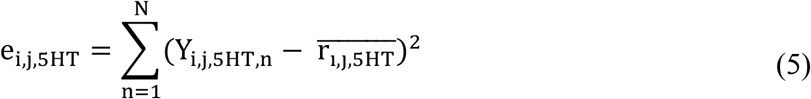

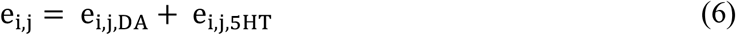

